# Recovery from anhydrobiosis in the tardigrade *Paramacrobiotus experimentalis*: better to be young than old and in a group than alone

**DOI:** 10.1101/2023.05.22.541721

**Authors:** Amit Kumar Nagwani, Iwona Melosik, Łukasz Kaczmarek, Hanna Kmita

**Author notes:** The authors have an equal contribution (Amit Kumar Nagwani, Iwona Melosik).

## Abstract

Desiccation-tolerant organisms can survive dehydration in a state of anhydrobiosis. Tardigrades can recover from anhydrobiosis at any life stage and are considered among the toughest animals on Earth. However, the factors that influence recovery from anhydrobiosis are not well understood. The study aimed to evaluate the effect of sex, age, the presence of other individuals and the combination of the number and duration of anhydrobiosis episodes on the recovery of *Paramacrobiotus experimentalis*. The activity of 1,200 individuals for up to 48 hours after rehydration was evaluated using ANOVA. Age was the main factor influencing return to activity, followed by the combination of number and duration of anhydrobiosis episodes, influence of the presence of other individuals, and sex. More individuals returned to activity after repeated short than repeated long anhydrobiosis episodes and older individuals were less likely to recover than younger individuals. In addition, when compared to single animals, the presence of other individuals resulted in higher number of active animals after dehydration and rehydration. The effect of sex was significant, but there was no general tendency for one sex to recover from anhydrobiosis better than the other one. The results contribute to a better understanding of the anhydrobiosis ability of *Pam. experimentalis* and provide background for full explanation of molecular, cellular and environmental mechanisms of anhydrobiosis.

## INTRODUCTION

The ability of some organisms to survive dehydration, resulting in almost complete loss of body water (desiccation) and entering a state of reversible suspension, is called “anhydrobiosis”, which comes from the Greek for “life without water” and indicates “desiccation tolerance” [1–5]. Anhydrobiosis is extremely important for survival in harsh environments with periodically unavailable water, which can affect growth and reproduction. It also affects lifespan and may therefore slow down the rate of evolution [4,6].

However, it has been shown that the longer the time spent in anhydrobiosis, the longer it takes to return to activity. Exceeding a certain critical period of desiccation can lead to the death of the organism [7–9]. As water availability is one of the most important factors for life, a full understanding of the underlying mechanisms of anhydrobiosis is crucial for the development of technologies based on organism tolerance to desiccation. The discovery and understanding of these mechanisms could have an impact on several areas of research, including DNA protection and repair mechanisms, the preservation of biological materials for clinical applications or food production, and enzymes working in a small amount of water [e.g., 4, 10, 11].

Tardigrades (commonly called water bears) are an important group of invertebrates due to their place between two major invertebrate model organisms, i.e., *Caenorhabditis elegans* and *Drosophila melanogaster*, and their aquatic-to-terrestrial transition, which may provide insight into the evolution of mechanisms that allow adaptation to stressful conditions [2]. Like other invertebrates, such as nematodes and rotifers, tardigrades show a remarkable ability to survive in an anhydrobiotic state at all life stages [e.g., 13–15], although not with the same success for all species and life stages [16, 17].

Tardigrades can serve as an excellent model in biological research, including for the impact of phenotypic and environmental factors on anhydrobiosis as a survival strategy. Indeed, various aspects of tardigrade biology have already been intensively studied, including reproduction [e.g., 18, 19], dormancy strategies [e.g., 2, 20–22], mechanisms of adaptation to the most extreme environments [e.g., 23, 24], phylogenetic relationships [e.g., 25–27], metabolic functions [e.g., 28] and experience of exposure to space conditions [e.g., 11, 29].

Anhydrobiosis in tardigrades is a complex phenomenon. It includes entering, permanent, and leaving steps, which correspond to dehydration (i.e., a tun formation), tun state (i.e., desiccated state) and rehydration, respectively [2]. These steps are fully elucidated at the level of the organism’s morphology [3, 13, 14, 20, 21], but full access to the underlying mechanisms requires consideration of additional factors. It is known that the survival rate of anhydrobiosis can be affected by the type of environment, feeding behaviour (e.g., diet), environmental/culture conditions (e.g., ambient temperature, water quality, culture substratum) and other factors such as overall body size, conditions of dehydration, as well as, the number, and duration of anhydrobiosis episodes [e.g., 9, 17, 30–33].

Up to now, *ca.* 1400 tardigrade species have been described [34], but fewer than 1.5% (*ca.* 20 species) have been studied for anhydrobiotic ability. Most of the studies have been mainly performed on parthenogenetic species. However, many bisexual species have been reported in tardigrades [e.g., 19]. In three bisexual species, females were predominantly analysed, or the sex was not specified [9, 35–36]. However, males can also occur in parthenogenetic lineages, suggesting a switch in the reproductive mode between parthenogenesis and bisexual reproduction [19, 37], which makes interpreting results even more complicated.

Some of the factors we examined that affect anhydrobiosis survival have already been evaluated, while others have not, or knowledge of them is incomplete, but they could potentially increase the risk of anhydrobiosis failure. Although individual age is generally considered to be a factor influencing anhydrobiosis survival [e.g., 30], the only study addressing this issue found no significant effect of age [31]. The duration of anhydrobiosis episodes is a known factor affecting survival. It is explained by cellular damages, the severity of which correlates with the duration of the tun state [e.g., 25, 35, 38]. The negative impact of repeated anhydrobiosis episodes on survival has also been shown [e.g., 35]. It has been hypothesized that the repeated entering and leaving steps cause additional cellular damage, which is supposed to be eliminated by feeding before the next episode of anhydrobiosis, thereby enhancing cellular repair mechanisms [30–31, 35, 39]. However, a comparative analysis of the combined effect of the number and duration of anhydrobiosis episodes has not been performed. In addition, the effect of sex and occurrence in the presence of other individuals on the return to activity has not yet been assessed.

In response to the shortcomings of research on anhydrobiosis in tardigrades, active individuals per test unit were examined after experimentally induced repeated anhydrobiosis episodes for the bisexual species *Paramacrobiotus experimentalis* [40]. The effect of the following factors was assessed: (1) sex, (2) age, (3) presence of other individuals termed here shortly “group influence” and (4) combination of the number and duration of anhydrobiosis episodes. In addition, the interactions between these factors were analysed. The approach used here differs from previous anhydrobiosis studies in several points because (1) males and females were analysed separately; (2) dehydration and rehydration were performed for single individuals and in the presence of other individuals, i.e., in groups; (3) individuals were divided into five age classes and (4) different number and duration of anhydrobiosis episodes were applied.

Results obtained are important for developing a better understanding of the process of anhydrobiosis, providing the background for describing underlying molecular, cellular and environmental mechanisms and their response to environmental stress.

## MATERIALS AND METHODS

### Cultures of Paramacrobiotus experimentalis

Females and males of *Pam. experimentalis* [40], were cultured together in covered, vented plastic Petri dishes (55 mm in diameter), with the bottom scratched with sandpaper to allow the animals to move. Individuals were coated with a thin layer of the culture medium, a mixture of spring water (Żywiec Zdrój S.A., Poland), and ddH_2_O in a 1:3 ratio. The culture medium was changed every week, and animals were fed with the rotifer *Lecane inermis* (strain 1.A2.15). provided by Dr Edyta Fiałkowska (Institute of Environmental Sciences, Jagiellonian University, Krakow, Poland). The Petri dishes were kept in the climate chamber POL EKO KK 115 TOP+ at 20°C, in the dark (24h), and with relative humidity (RH) of 40%.

On the basis of life history traits, five age classes, defined as growing, young, mature, late and old adults, were distinguished (Supplementary file, Table S1). They represent the following age ranges in days: 60–90, 120–150, 150–180, 240–270 and >300, and correspond to age classes 1-5, respectively. For the selected age classes an approximate ratio of 2:1 females to males was defined.

### Protocol for repeated episodes of anhydrobiosis

The protocol includes dehydration and rehydration procedure optimized for *Par. experimentalis* [33]. All experiments were performed in covered, vented plastic Petri dishes of 35 mm diameter lined at the bottom with white filter paper (grammage 85–87, Chemland Company, Poland). Females and males representing the distinguished age classes (Supplementary file, Table S1) were transferred using an automatic pipette into dishes filled with 450 µl of the culture medium. The dishes were placed into the climate chamber PolLab Q-Cell 140, and the individuals were allowed to dry slowly at 20 °C, with 40–50% RH, and in the dark for 72□h. The tun formation was checked once every 24 h by a brief 1-minute observation under the stereomicroscope (Supplementary file, Figure S1). After the tun formation, these conditions were maintained for 3 days or 30 days. Next the individuals were rehydrated and their return to activity was observed after 2h, 6h, 24h, and 48h. The rehydration was achieved by adding 3 ml of the culture medium to each Petri dish. Tuns were transferred using an automatic pipette to small glass cubes and kept at 20□°C and 40–50% RH, with light conditions regulated by seasonal changes in the day/night cycle (according to our observations, the photoperiod does not affect the return of *Pam. experimentalis* tuns to the active state).

Those individuals that returned to activity (defined here as coordinated movements of the body and legs, i.e., the onset of crawling) 48h after rehydration were subjected to another anhydrobiosis episode. The number of active individuals after each anhydrobiosis episode and at a given observation time (Supplementary file, Table S2) was used to calculate the activity score defined as the number of active individuals per test unit (3×10 individuals). All variants of anhydrobiosis applied in this study are summarised in Figure 1. A break of three days was allowed between the consecutive anhydrobiosis episodes, and the animals were fed. The feeding took place three days before the start of the next anhydrobiosis episode. Females and males of different ages were subjected to repeated anhydrobiosis in the presence of other individuals, i.e., in groups (10 specimens per Petri dish) or singly (one individual in each of 10 Petri dishes). In total of 1,200 individuals were analysed, 600 each in short and long anhydrobiosis episodes with the same duration of dehydration and rehydration steps (see Supplementary file, Table S2 for more details).

**Figure 1.**
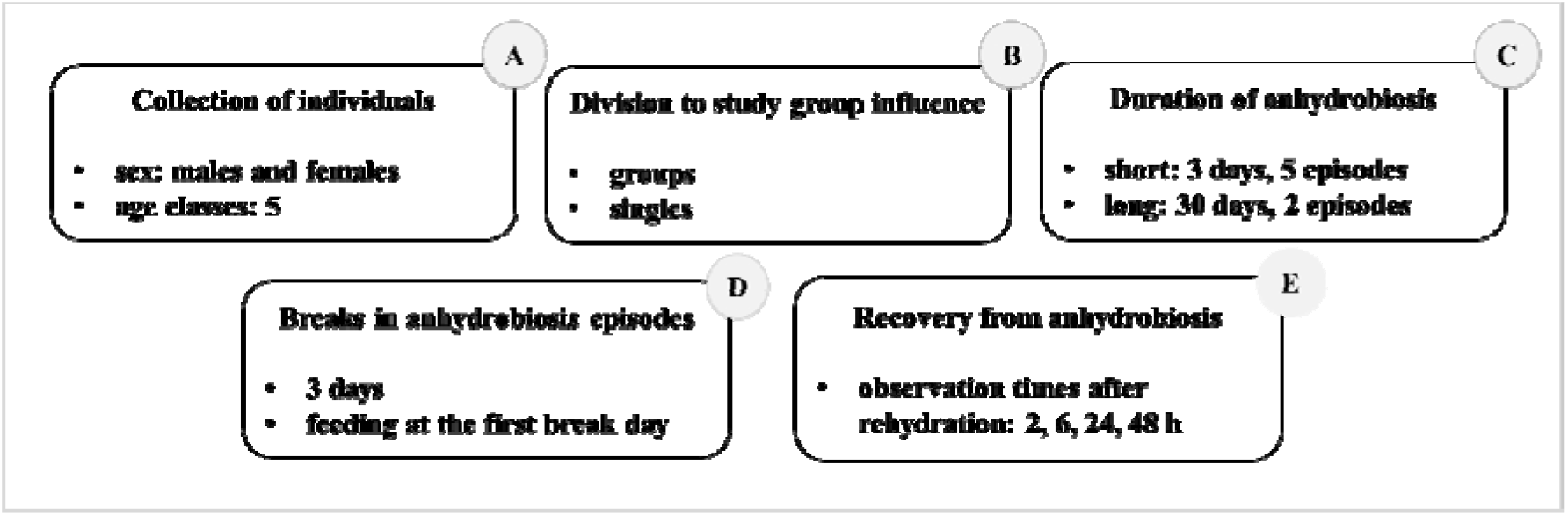
Graphic representation of the repeated anhydrobiosis experiment performed in *Paramacrobiotus experimenatlis.* A-E, the applied steps of the experiments. The term “anhydrobiosis duration” refers to combination of the number and duration of anhydrobiosis episodes.

### Statistical analysis

A multivariate repeated measures analysis of variance (RM_ANOVA) was performed using the GLM procedure to compare differences in the activity of individuals after rehydration according to the levels of the main order factors considered, i.e. the combined effect of the number and duration of anhydrobiosis episodes, the group effect, the age and sex of the individuals, and the interaction between these factors. Details of the data transformation and statistical tests used to assess the ANOVA assumptions are in Supplementary file. Eta squared, an integral part of ANOVA, was used to assess the size of one or more effects (i.e., the proportion of variance accounted for by the effects). After each anhydrobiosis episode for a given observation time (2h-48h), the number of active individuals in each sample unit was treated as a repeated measure. Pairwise comparisons were evaluated using Tukey’s post hoc test when the F ratio was statistically significant (at alpha <0.05) for the main determinants and their interactions [41]. The effect of the number of episodes applied for long and short anhydrobioses in the context of active individuals after rehydration was evaluated using the Student’s t-test or the one-way ANOVA. All statistical analyses were performed in Statistica version 13.0 (StatSoft, Poland).

## RESULTS

### General remarks

The number of active individuals of *Pam. experimentalis* (see Suplementary file, Table S2 for raw data) depended most strongly on the observation time after rehydration and the age of the individuals (Eta-squared = 0.633 and 0.553, *p*=0.001, respectively). The effect of the combination of the number and duration of anhydrobiosis episodes, and the presence of other individuals (i.e., group influence) appeared to be of medium size (Eta-squared = 0.133 and =0.103, respectively, *p*=0.001). In contrast, the proportion of variance accounted for by sex was small but significant (Eta-squared = 0.043, *p*=0.05) (Supplementary file, Table S3).

The most significant interactions were between observation time, age of individuals, and the combination of the number and duration of anhydrobiosis episodes (Eta-squared = 0.260, *p*=0.001, or interactions of two of these variables (Figure 2; Supplementary file, Table S3). Some of the other interactions were also significant. However, Eta-square indicates that the proportion of variance a given interaction can explain was relatively small (Eta-square from 0.02 to 0.09).

**Figure 2.**
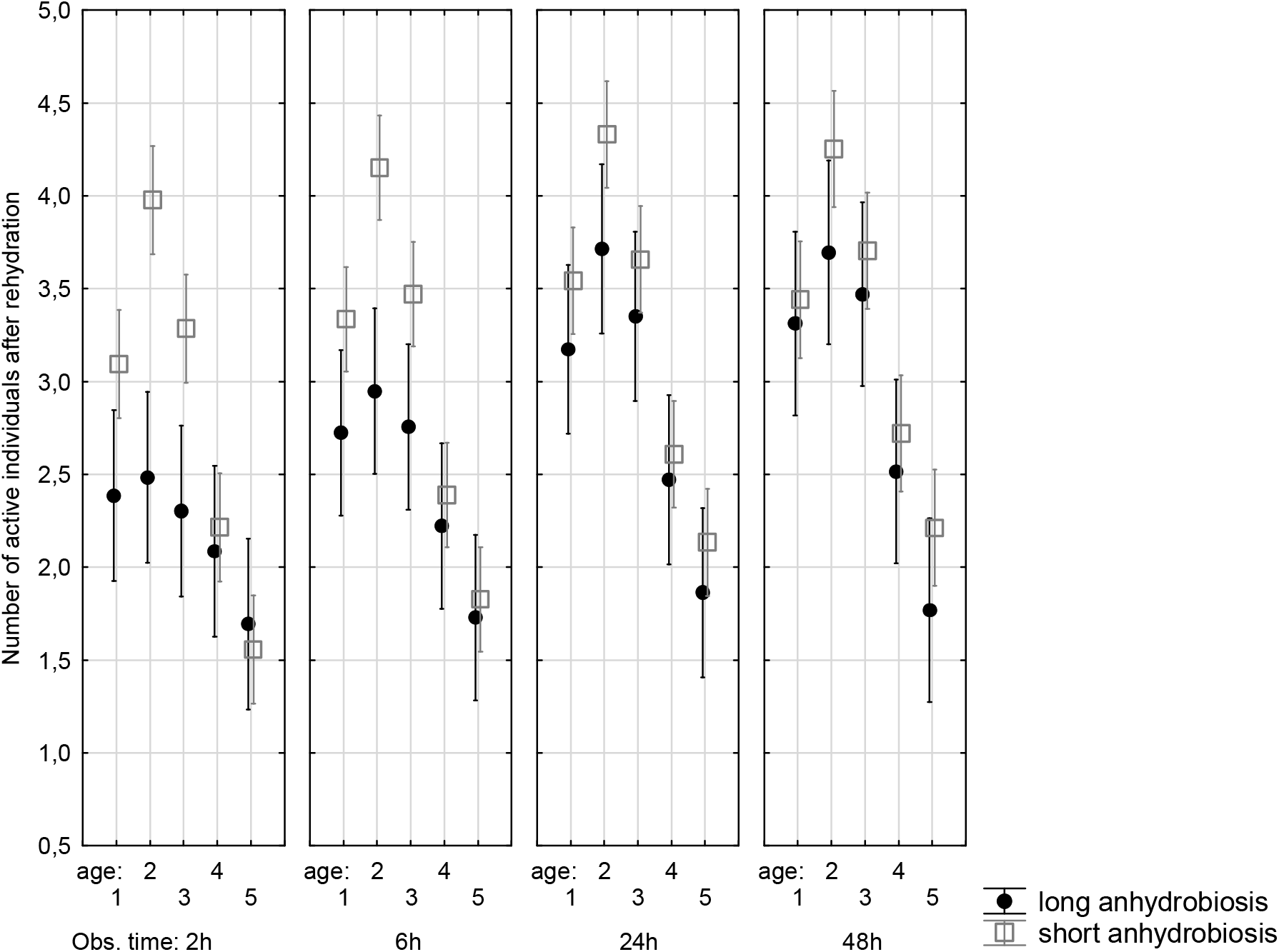
Number of active individuals of *Paramacrobiotus experimentalis* observed at 2h-48h after rehydration, taking into account their age and the combined effect of the number and duration of anhydrobiosis episodes (long and short anhydrobiosis). Age (1-5); the selected age classes; Obs. time - observation time. Expected marginal means and 0.95% confidence intervals are presented.

### Effect of age

Based on the post hoc tests, more individuals were active after rehydration in the younger age classes (age classes 1–3) than in the older age classes (age classes 4–5), regardless of the combination of number and duration of anhydrobiosis episodes (Figure 2). Taking into account the mean values, the younger age classes (1-3) had, on average, 37% more active individuals after rehydration than the older age classes (4-5) (Supplementary file, Table S4).

Statistically significant differences in the number of active individuals after repeated long and short anhydrobiosis were observed especially at the initial observation time after rehydration (2h) and mainly for individuals representing the younger age classes (Figure 2 and post hoc tests (not shown)). The differences diminished with the time of observation and became insignificant at 48h after rehydration. Based on the mean values at 48h after rehydration, the young adults (age class 2) returned to activity 52% and 49% more successfully than the old adults (age class 5) after repeated long and short anhydrobiosis, respectively (Supplementary file, Table S4). The growing adults (age class 1) returned to activity 10% and 19% slower, the mature adults (age group 3) 7% and 14% slower, and the late adults (age class 4) 32% and 37% slower than individuals representing the young adults (age class 2) (Supplementary file, Table S4).

### Effect of combination of the number and duration of anhydrobiosis episodes

The number of active individuals decreased significantly with the increasing number and duration of anhydrobiosis episodes but increased with the observation time after rehydration (Figure 2). In general, significant positive correlations (*p*<0.05) were found between the combined effect of the number and duration of anhydrobiosis episodes and observation at 2h and 6h after rehydration. Although still positive, these correlations became statistically insignificant as rehydration time progressed (observations at 24–48h; Supplementary file, Table S5).

In the subsequent analysis, two episodes of long anhydrobiosis were compared with the first two episodes of short anhydrobiosis to distinguish the effect of the duration of anhydrobiosis from the number of anhydrobiosis episodes (2 *vs*. 5). With a 90% difference in the duration of the tun state (i.e., 6 *vs*. 60 days), 25% to 36% more active individuals were observed after short anhydrobiosis than after long anhydrobiosis, depending on the observation time (Supplementary file, Table S6). Based on post hoc tests, significant differences were found between short and long anhydrobiosis in the number of active individuals after rehydration at each observation time (2–48h; one-way ANOVA, Wilk’s Lambda = 0.544, (F (4,75) = 15.728, *p*<0.001), Figure 3).

**Figure 3.**
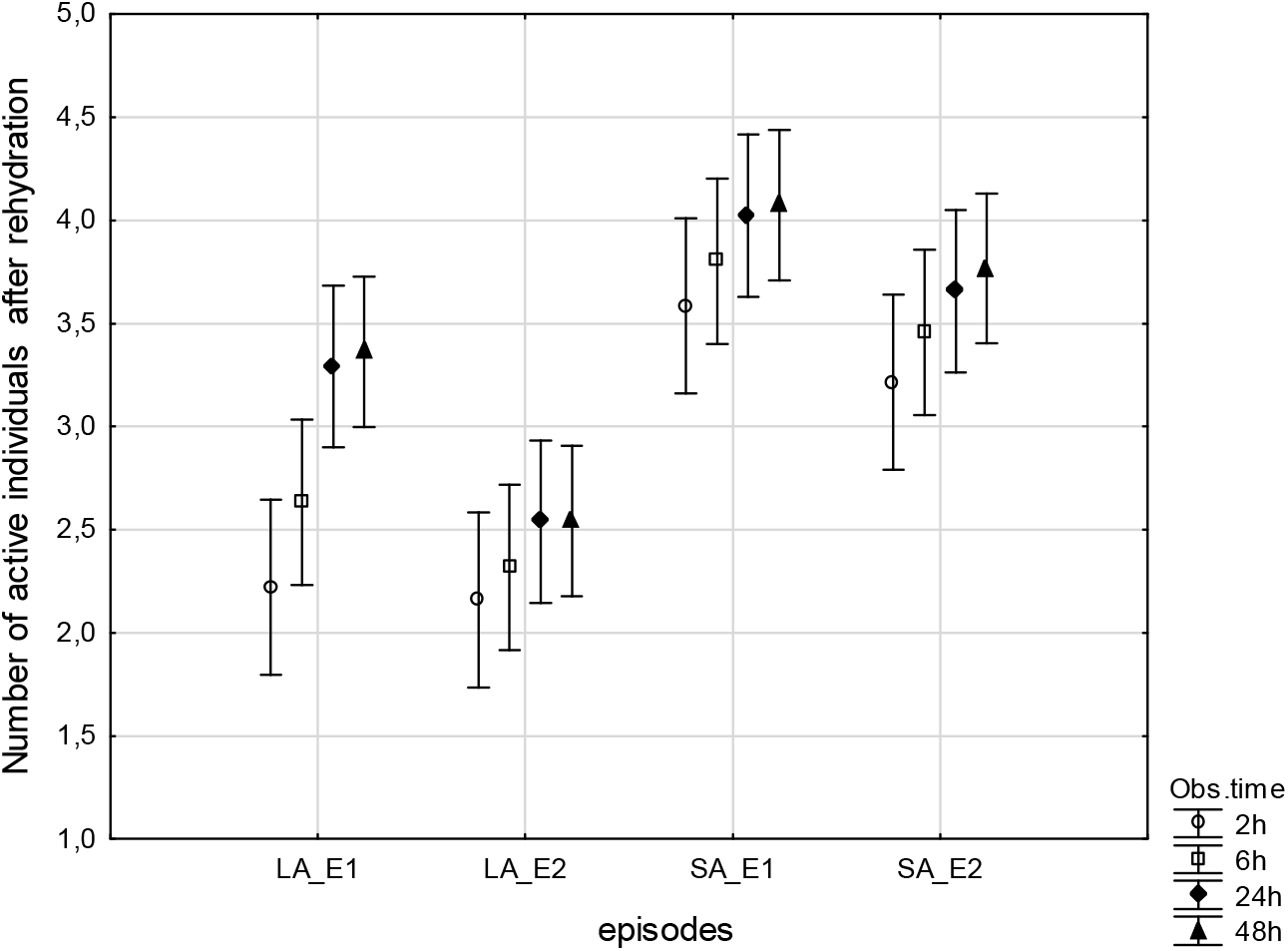
Number of active individuals of *Paramacrobiotus experimentalis* observed at 2h-48h after rehydration following the first two episodes of short and long anhydrobiosis. LA_E1 and LA_E2, the first and second episode of long anhydrobiosis; SA_E1 and SA_E2, the first and second episode of short anhydrobiosis; Obs. time - observation time.

The comparison between consecutive short anhydrobiosis episodes showed no statistically significant differences in the number of active individuals, except a significant difference at 48h after rehydration between the fourth and fifth episode.

### Effect of the presence of other individuals

The presence of other individuals (i.e., group influence) during anhydrobiosis significantly influenced the number of active individuals after rehydration (*p*<0.001, Supplementary file, Table S3). Furthermore, interactions between observation time after rehydration, the combination of the number and duration of anhydrobiosis episodes, and the group influence or interactions of two of these specified factors were significant (Supplementary file, Table S7). Irrespective of observation time and combination of the number and duration of anhydrobiosis episodes, more individuals were active in groups after rehydration when compared to single animals (Figure 4). Based on the mean values, the number of active individuals experiencing repeated long anhydrobiosis individually was 20% lower than in groups, and for short anhydrobiosis, the difference between groups and individuals was 6%. However, the Eta-squared indicates that the proportion of variance explained by the interaction was low (Eta-squared equal to 0.03).

**Figure 4.**
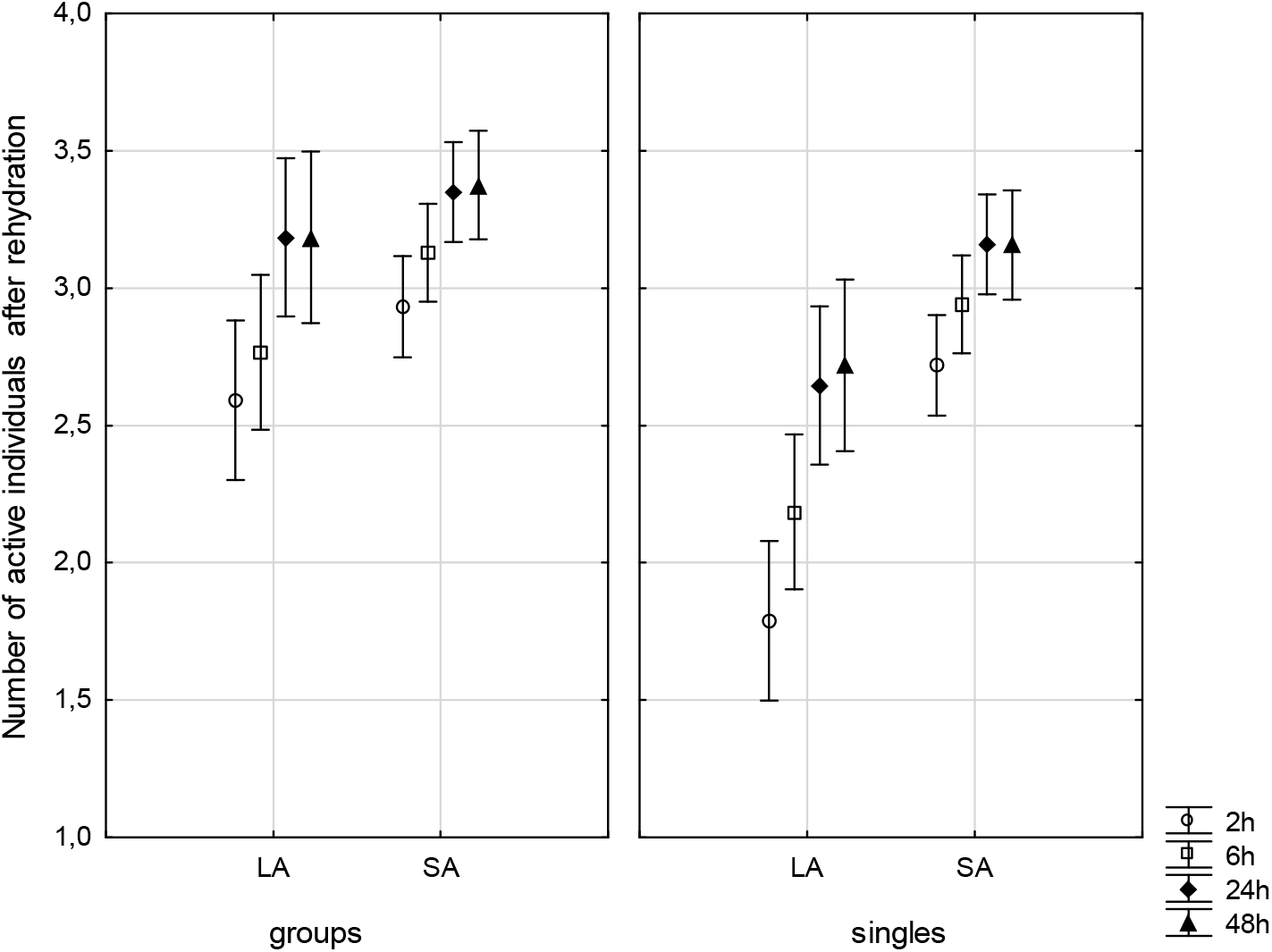
Number of active individuals of *Paramacrobiotus experimentalis* according to the combined effect of number and duration of anhydrobiosis episodes, and observation times after rehydration (2h-48h; F(3, 300)=2.687, *p*=0.047). LA and SA, long and short anhydrobiosis, respectively; Obs. time - observation time. Expected marginal means and 95% confidence intervals are shown.

### Effect of sex

The main effect of sex was statistically significant (*p*<0.05; Supplementary file, Table S3). However, there was no general trend to conclude that one sex had a higher number of active individuals after rehydration than the other one. Significant differences between females and males in the number of active individuals were only observed at 2h and 6h after rehydration for the repeated short and long anhydrobiosis, respectively. We also found no significant differences between the sexes when analysing the different age classes, with one exception concerning short anhydrobiosis for the oldest age class at 2h after rehydration (Figure 5).

**Figure 5.**
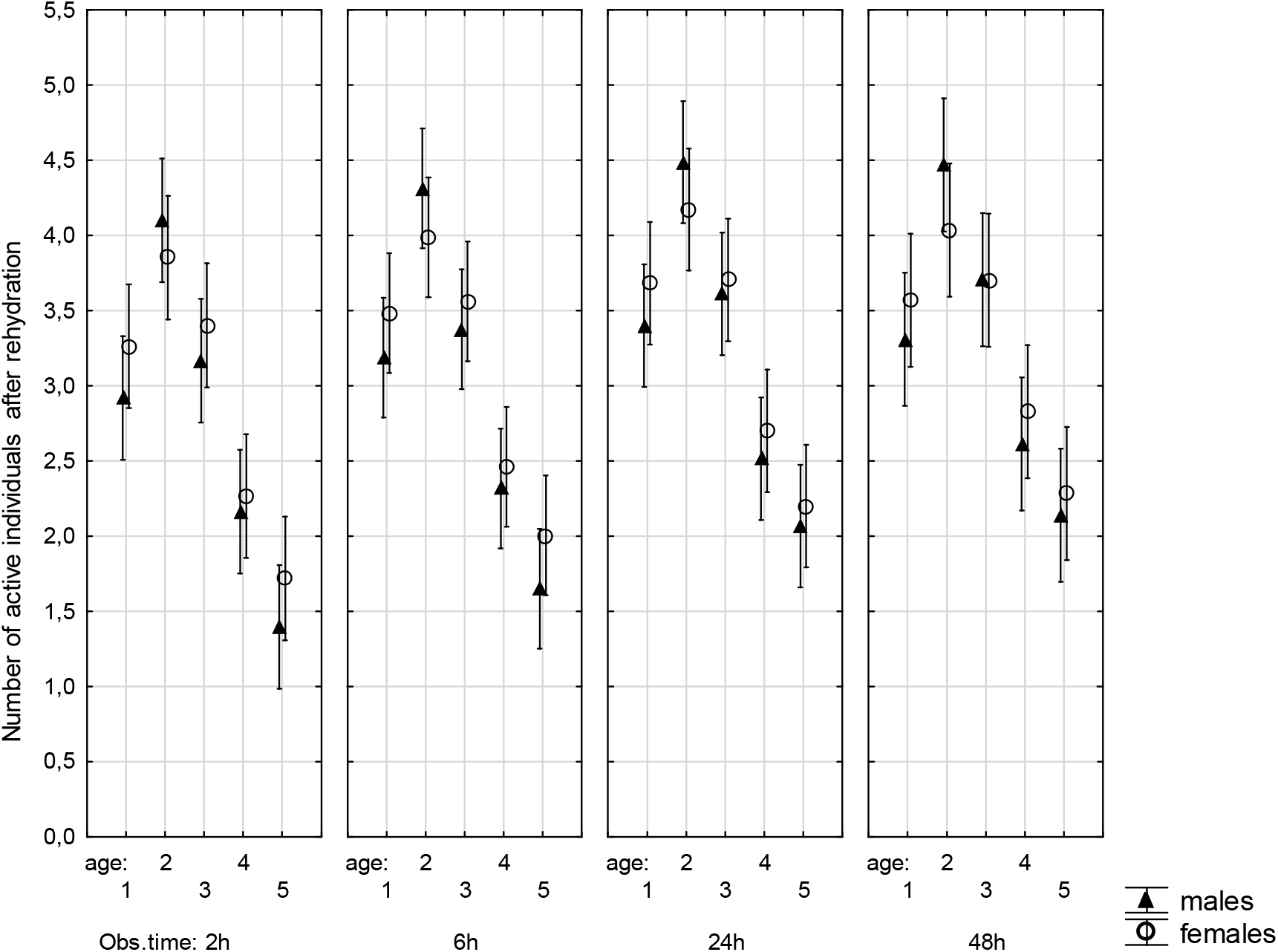
Number of active individuals of *Paramacrobiotus experimentalis* in the context of their sex and age, as well as observation time after rehydration. Age (1-5); the selected age classes; Obs. time; observation times. Expected marginal means and 95% confidence intervals are shown; see the Materials and methods section for details. Obs. time - observation time.

The interaction between observation time, age and sex of individuals appeared to be significant (F(12, 300) = 2.537, *p*<0.05), meaning that both females and males differed in the number of active individuals between older and younger age classes, particularly at the later observation time (24–48h) after rehydration. Other interactions where sex was one of the main factors were not significant (Supplementary file, Table S3). Mean numbers of active individuals representing females and males in the context of age, combination of the number and duration of anhydrobiosis episodes and observation time after rehydration (2–48h) are presented in Supplementary file, Table S8.

## DISCUSSION

The present study demonstrates the potential of the newly-described tardigrade species *Pam. experimentalis* to return to activity after repeated short and long anhydrobiosis. This potential was investigated by the determination of the activity of individuals after rehydration (defined here as coordinated movements of a body and legs, i.e., the onset of crawling) concerning their age, sex, and whether they occurred in the presence of other individuals (i.e., in a group) or not (i.e., single) during anhydrobiosis steps. The study also focuses on the combined effect of the number and duration of anhydrobiosis episodes. Some of these factors have not been studied before (e.g., the effect of sex) or only limited knowledge of them was previously available. The impact of these factors was evaluated to guide future research on tardigrades anhydrobiosis and to provide a more comprehensive characterisation of the species under study. In a broader sense, this study may also contribute to the validation of two hypotheses proposed to explain anhydrobiosis effect on aging, i.e., ”the Sleeping Beauty” or” the Picture of Dorian Gray” [4, 14, 17, 42].

The main findings of the study can be summarised as follows: (1) recovery from anhydrobiosis declined with age; (2) regardless of age, the number of active individuals decreased significantly with increasing number and duration of anhydrobiosis episodes, but increased with observation time after rehydration; (3) some individuals were able to return to activity after five short or two long anhydrobiosis episodes; (4) individuals in groups returned to activity after rehydration more efficiently than those treated individually; (5) sex appeared to be a significant predictor of the number of active individuals after rehydration, but the effect size was very small. The results indicate that three of the most important factors influencing return to activity after anhydrobiosis are age, combination of the number and duration of anhydrobiosis episodes and the group influence (i.e., the influence of other individuals’ presence). These factors could be considered in studies of anhydrobiosis in tardigrades The impact of sex on return to activity after rehydration requires further research.

The lifespan of tardigrades varies between species, populations and individuals, and ranges from a few weeks to two years, not counting the time spent in the dormant state [17, 43–45]. Accordingly, the most obvious finding to emerge from the analysis performed is that young adults of *Pam. experimentalis* (age in days 120–150) showed the best recovery from repeated anhydrobiosis. In contrast, the recovery was the worst for the oldest individuals (age in days >300). This finding supports the results of other studies linking the age of individuals with their ability to recover from anhydrobiosis. In *Milnesium tardigradum* higher recovery rates after anhydrobiosis were observed for younger individuals (from 37 to 149 days old) than for older ones (163 to 191 days old) [31]. Similarly, studies on the nematode *Panagrolaimus rigidus* showed that increasing age had a negative effect on recovery from anhydrobiosis [42]. However, the ability of embryos to survive anhydrobiosis increased with age in this species [46]. In the population of the *Richtersius coronifer* from Sweden, the body size affected the survival of anhydrobiosis, i.e., larger individuals showed a lower probability of return to activity than medium-sized ones [30]. Assuming a correlation between body size and age [47], younger individuals would recover from anhydrobiosis better than older ones, but this may apply to limited age ranges. Accordingly, in our experiments, young adults (age in days 120–150) showed a better return to activity after rehydration than growing adults (age in days 60–90). Thus, our results are somewhat consistent with data for the bdelloid rotifer *Macrotrachela quadricornifera*, for which adult individuals showed better recovery from anhydrobiosis than juveniles and eggs [48, 49].

Among the tardigrades tested, ability of anhydrobiosis varies over a wide range [9, 14, 17, 33]. This may be related to abiotic factors, such as the moisture content in the natural environment, the innate ability of the animal to recover and specific conditions for entry into anhydrobiosis and rehydration [7]. Species living in constantly moist habitats tend to have a lower ability to tolerate drought by anhydrobiosis than those living in dry environments [11, 50]. It has been suggested that the upper limit of recovery from anhydrobiosis by tardigrades can be counted in years but does not exceed ten years [51]. However, other available data indicate that some tardigrade species can return to activity when the tun (desiccated) state lasts for up to 15–22 years [e.g., 52]. For semi-terrestrial tardigrades, also represented by *Pam. experimentalis*, the ability of anhydrobiosis has been studied, among others, for the eutardigrade *Ramazzottius oberhaeuseri* and the heterotardigrade *Echiniscus* spp. It was shown that under natural conditions *Ram. oberhaeuseri* recovered from the tun state lasting 1192 days with an average revival of 21.7% but could survive in this state for up to 1604 days. In *Echiniscus* spp., after 706 days of anhydrobiosis, the average revival was 9.9%, but the species could tolerate the tun state lasting for up to 1085 days [8].

Because of the different species tested and the methods used, our study cannot be directly compared with previous studies. Differences include, among others, the number and duration of anhydrobiosis episodes. In the case of *Pam. experimentalis*, significant differences in the number of active individuals after repeated anhydrobiosis persisted during the initial observation times after rehydration (2–6h), but became insignificant over time, indicating important differences in the rate of recovery. The observed return to activity at 48h after repeated short and long anhydrobioses was consistent with our previous experiments on *Pam. experimentalis* showing that this species has a high capacity for anhydrobiosis, as the average recovery rate of individuals after 240 days of tun state was 43% [33]. However, representatives of other populations of the species would need to be studied to verify correlations between the conditions of the natural environment and anhydrobiosis ability for this species. Previous studies are contradictory regarding population differences in tardigrades’ recovery from anhydrobiosis, which is explained by the intraspecific variation of physiology and/or habitat properties [6, 33, 53].

We found a significant positive correlation, at least for the initial observation time after rehydration, between return to activity and combination of the number and duration of anhydrobiosis episodes, confirming previous findings in nematodes [54] and various tardigrade species [e.g., 7, 14, 25, 33, 35, 38]. Namely, the longer the tun state, the more time the animals need to return to activity. We also showed that some individuals could recover after five repeated short or two repeated long anhydrobiosis episodes. This finding for *Pam. experimentalis* is generally consistent with that for the tardigrade *Mil. tardigradum* and *Ric. coronifer*, which can survive up to six and even nine consecutive short anhydrobiosis episodes, respectively [31 and 35]. For the three species, decreasing ability to form proper tuns was only observed for *Ric. coronifer* and the authors suggest that the decrease in anhydrobiotic performance could be explained by the lack of animal feeding between episodes [35]. Accordingly, the feeding was applied in the studies of *Mil. tardigradum* [31] and in our study of *Pam. experimentalis*. However, it cannot be excluded that other factors may contribute to the difference, including the time between anhydrobiosis episodes, their duration, studied species or the source of specimens, i.e. laboratory culture or environment. Nevertheless, the significant difference in the number of active individuals after rehydration between two long anhydrobiosis episodes and the lack of the difference between two short episodes supports the crucial influence of the duration of the tun state on recovery from repeated anhydrobiosis. Accordingly, it is generally accepted that the duration of the dry state is decisive for recovery from anhydrobiosis [e.g., 14], but it has also been proposed that recovery from anhydrobiosis may be influenced by factors acting during dehydration and rehydration [55], making the distinction between the effects of the number and duration of anhydrobiosis episodes more complex.

The effect of the presence of individuals in a group on recovery from anhydrobiosis has not been widely studied although it has been shown to be related to aggregations [56]. Namely, it has been shown in the tardigrade *Ric. coronifer* that aggregations of individuals can improve the survival of anhydrobiosis because may contribute to a reduction in the body surface area exposed to desiccation and, thus, to a reduction in the rate of water evaporation, thereby increasing the chance of return to activity. This is assumed to be of a crucial meaning for survival of rapid desiccation [56]. From an ecological point of view, the positive consequences of individuals aggregation and its role in animal recovery was highlighted by Jönsson [6], who noted that population density could promote aggregation and *vice versa*. This might explain, for example, the distribution and abundance of tardigrade species in xerothermic habitats [6].

The performed studies showed that tardigrades in groups recovered from anhydrobiosis better than single individuals. However, we did not observe typical aggregations of individuals (Supplementary file, Figure S1). Therefore, it appears that under certain conditions, formation of aggregates is not indispensable for successful anhydrobiosis. The decisive factor maybe be the amount of water applied during tun formation and/or the rate of dehydration [50]. We can also speculate that the presence of other individuals could be a source of chemical signals that could be released in response to dehydration and/or rehydration, and enhance *Pam. experimentalis* recovery from anhydrobiosis. Neither the nature of the signals nor their relationship to the age of individuals is known. Nevertheless, research on the mating behaviour of tardigrades illustrate the role of chemical communication between tardigrades [57–58]. On the other hand, the absence of the aggregates may explain the lower return to activity after the first episode of long anhydrobiosis (30 days) when compared with the relevant data available for *Pam. experimentalis* [33], although the deterioration was not observed for the first episode of short anhydrobiosis (3 days). However, the age and sex were not considered in that study, and well as the shortest duration of the tun state was 7 days, that does not allow for deeper comparison.

Although the effect of sex was statistically significant, a general trend for females to recover better from anhydrobiosis than males was not observed. This aspect requires further research because cannot be simply correlated with the calculated 2:1 female-biased sex ratio. Similar value of the female-biased sex ratio (2:1) was reported for *Paramacrobiotus sp.* TYO [57]. In both species the ratio was determined for animals not undergoing anhydrobiosis and in the case of *Pam. experimentalis* in all age classes. Additionally, we observed only a marginally lower maximum lifespan for *Pam. experimentalis* males (400 days) when compared to females (420 days). The female-biased sex ratio is an inevitable issue for considerations on the evolution of sexual reproduction that still remains a fascinating enigma in biology. It is usually assumed that males are costly, and their cost can be reduced by decreasing the ratio of males to females [59]. This appears to be a rule in tardigrades. Accordingly, although sex ratio close to 1:1 was noted in tardigrades of the genus *Macrobiotus* C.A.S. [60], in *Ramazzottius* sp. the equal sex ratio was only found in a limited number of samples, and generally, a female-biased sex ratio was observed [61].

## CONCLUSIONS

The most significant predictor of recovery from repeated anhydrobiosis in *Pam. experimentalis* is the age of the individual. The combination of number and duration of anhydrobiosis episodes, and the presence of other individuals are secondary predictors. Although there was a little evidence for the effect of sex on the recovery, this factor should be further analyzed in different populations of this species and other bisexual tardigrade species.

The analyzed predictors may help to understand the molecular and cellular mechanisms governing tardigrade anhydrobiosis and the response of these mechanisms to environmental stress. This research may also be helpful in the context of evolutionary adaptations and responses to droughts caused by climate change and water shortage.

## Acknowledgment

The study was supported by the research grants of the National Science Centre, Poland, NCN 2016/21/B/NZ4/00131 and 2021/41/N/NZ3/01165, and partially conducted in the framework of activities of BARg (Biodiversity and Astrobiology Research Group). Amit Kumar Nagwani is a scholarship passport holder of the Interdisciplinary Doctoral Studies at the Faculty of Biology, Adam Mickiewicz University, POWR.03.02.00-00-I006/17. The authors would also like to thank Cambridge Proofreading LLC (http://proofreading.org) for their linguistic assistance. Technical support of Tomasz Bartylak and Pushpalata Kayastha is highly appreciated.

## Author contributions

Conceptualization, H.K. and A.K.N.; data curation, A.K.N and I.M; investigation, A.K.N. and I.M.; methodology, Ł.K., H.K. and I.M.; statistical analysis, I.M.; validation, I.M., H.K., and Ł.K.; supervision, H.K. and Ł.K.; writing, I.M., H.K., Ł.K., and A.K.N. All authors accepted the final version of the manuscript.

## Data Availability Statement

All relevant data are within the paper and the Supplementary file.

## Ethical approval

Samples of *Pam. experimentalis* were collected according to research permission from the Direction Generale des Forests, Direction de la Valorisation des Ressources Forestieres, Antananarivo, Madagascar (autorisations de recherche: No: 260/15-MEEMF/SG/DGF/DCAP/SCBT and Service de la Gestion Faune et Flore No: 056N-EA03/MG18).

