## Supplementary file for "Recovery from anhydrobiosis in the tardigrade *Paramacrobiotus experimentalis*: better to be young than old and in a group than alone"

*Cultures of Paramacrobiotus experimentalis*

Individuals of *Pam. experimentalis* were extracted in 2019 from moss samples collected in a tropical forest located in the Toamasina Province in Eastern Madagascar (18°56′37″S, 48°30′52″E, 717 m asl). The region is known for long periods of drought that recur annually, probably due to El Niño events [1]. The Latin name “*experimentalis*” refers to the species being easy to culture and valuable in different types of research, including studies on anhydrobiosis [2]. It is a bisexual species, and sex differences between males and females concern body size and primary sexual characteristics (unpaired internal genitalia). However, the ratio of males to females in a population (the sex ratio) for this species has not been determined yet.

The set of founder individuals was extracted from the moss sample according to the standard method [3]. In short, specimens were isolated by placing moss in a beaker filled with ddH_2_O up to 250 ml. After 6h, the moss was vigorously shaken with the use of metal tweezers. Then, the moss was squeezed over the beaker. The supernatant containing tardigrades as well as moss particles was stirred and poured into a 250 ml cylinder. After 30 min (when all particles fell onto the cylinder bottom), the top of ca. 150 ml of water was discarded. The remaining 50 ml was stirred and poured onto Petri dishes. Tardigrades found were extracted with a fine Pasteur pipette.

Females and males were reared together and the laid eggs were collected to prepare, and maintain a living assemblage of mixed-age animals. The assemblage was divided into groups that differed in age by a month. The collected eggs were used to form subsequent groups. The groups were kept in separate Petri dishes and characterized by selected life history traits (Table S1), including vitality rate (calculated as the ratio of active individuals and the total number of individuals, i.e., active and inactive), average total body length, and fertility (measured by the average number of eggs laid per female). Females and males were distinguished based on morphological characteristics, i.e., body shape (males are more slender than females), total body length (males are smaller than females), and the presence of eggs in the female’s ovary. The accuracy of sexes identification was confirmed by observing gonads as described in [4]. Total body length and the presence of eggs were assessed using a stereomicroscope (OLYMPUS SZ61). The calibrated grid of the stereomicroscope was used to measure the total body length of animals within a given area with an accuracy of 10 µm.

1. Gautier, L., Lowry II, P. P. & Goodman, S. M. in *The New Natural History of Madagascar* (ed. Goodman, S. M.) 452–464 (Princeton University Press: New Jersey) (2022).
2. Roszkowska, M. *et al.* Tips and tricks how to culture water bears: simple protocols for culturing eutardigrades (Tardigrada) under laboratory conditions. *Eur. Zool. J.* **88,** 449–465. <https://doi.org/10.1080/24750263.2021.1881631> (2021b).
3. Stec, D., Smolak, R., Kaczmarek, Ł. & Michalczyk, Ł. An integrative description of *Macrobiotus paulinae* sp. nov. (Tardigrada: Eutardigrada: Macrobiotidae: *hufelandi* group) from Kenya. *Zootaxa*, **4052,** 501–526 <https://doi.org/10.11646/zootaxa.4052.5.1> (2015).
4. Kaczmarek, Ł. *et al.* Integrative description of bisexual Paramacrobiotus experimentalis sp. nov. (Macrobiotidae) from republic of Madagascar (Africa) with microbiome analysis. *Mol. Phylogenet. Evol.* **145**, 106730. <https://doi.org/10.1016/j.ympev.2019.106730> (2020).

**Table S1**. Characterisation of age classes distinguished for *Paramacrobiotus experimentalis* based on the determined life history trait analysis. The vitality rate denotes the ratio of active individuals in the given age class and the total number of individuals in that class (i.e., active and inactive). Abbreviations: n, the sample size; ND, no data

| **Age classes** | **Age-range in days**  **(assigned group name)** | **Vitality rate**  **(± SD) [%]** | **Average body length**  **(± SD) [μm]** | | **Average number of laid eggs per female (± SD)** |
| --- | --- | --- | --- | --- | --- |
|  | | | **Females** | **Males** |  |
| 1 | 60-90  (growing adults) | 75.4 ± 6.7 | 530 ± 30  n = 25 | 430 ± 20  n = 25 | 123 ± 12  n = 100 |
| 2 | 120-150  (young adults) | 79.0 ± 2.4 | 750 ± 30  n = 25 | 570 ± 50  n = 25 | 142 ± 21  n = 80 |
| 3 | 150-180  (mature adults) | 81.4 ± 3.8 | 750 ± 40  n = 20 | 590 ± 50  n = 20 | 162 ± 32  n = 70 |
| 4 | 240-270  (late-age adults) | 52.1 ± 0.3 | 750 ± 30  n = 15 | 600 ± 30  n = 15 | 60 ± 8  n = 40 |
| 5 | >300  (old adults) | 40.3 ± 0.5 | 750 ± 30  n = 10 | 600 ± 30  n = 10 | ND |

**Table S2**. Number of *Paramacrobiotus* *experimentalis* individuals that returned to activity after each of anhydrobiosis episodes. Individuals that returned to activity after the previous episode were used in the next one. Each variant of the experiment included 3 replicates of 10 individuals at its beginning.

Females: five episodes of short anhydrobiosis

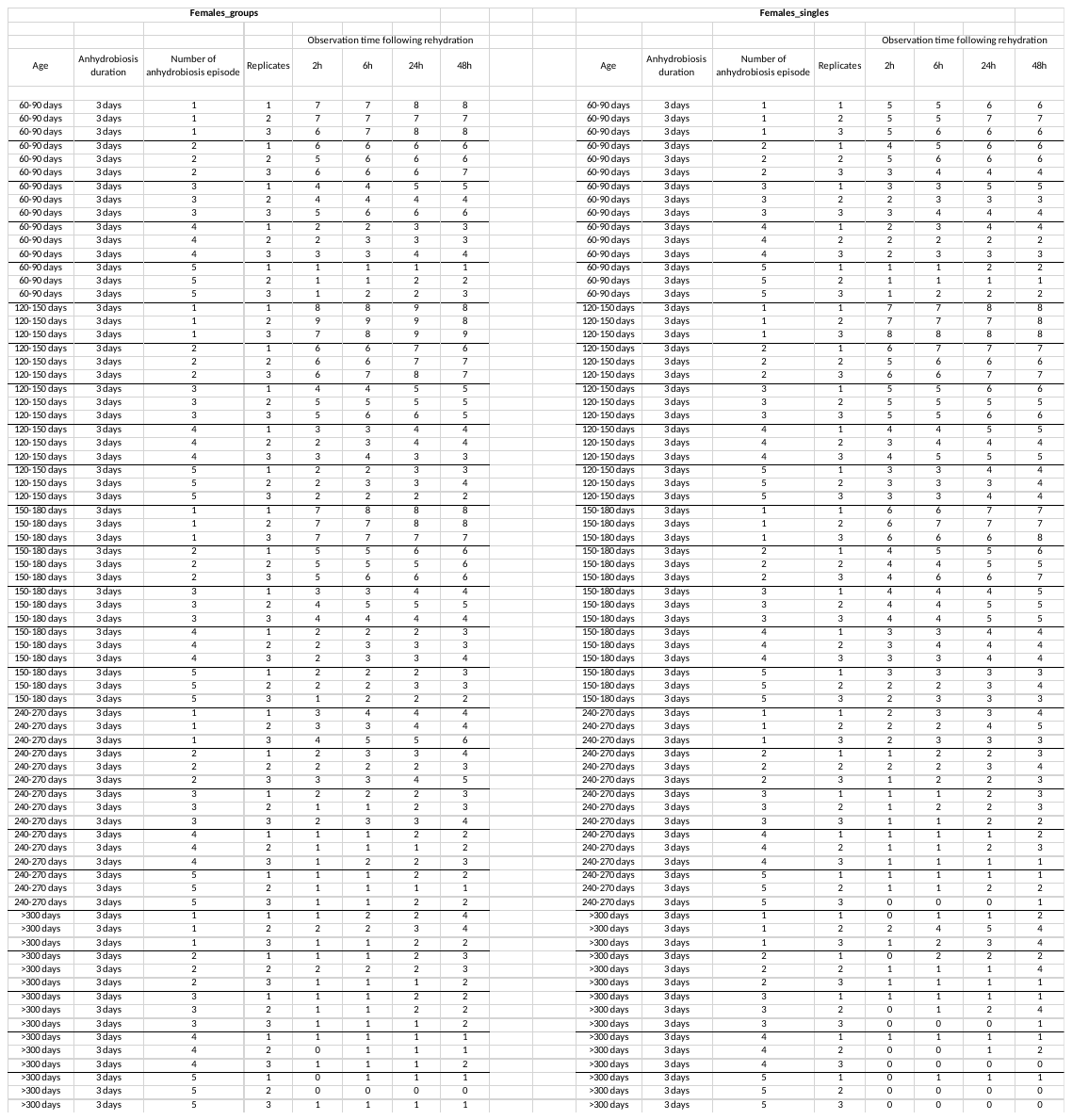

Males: five episodes of short anhydrobiosis

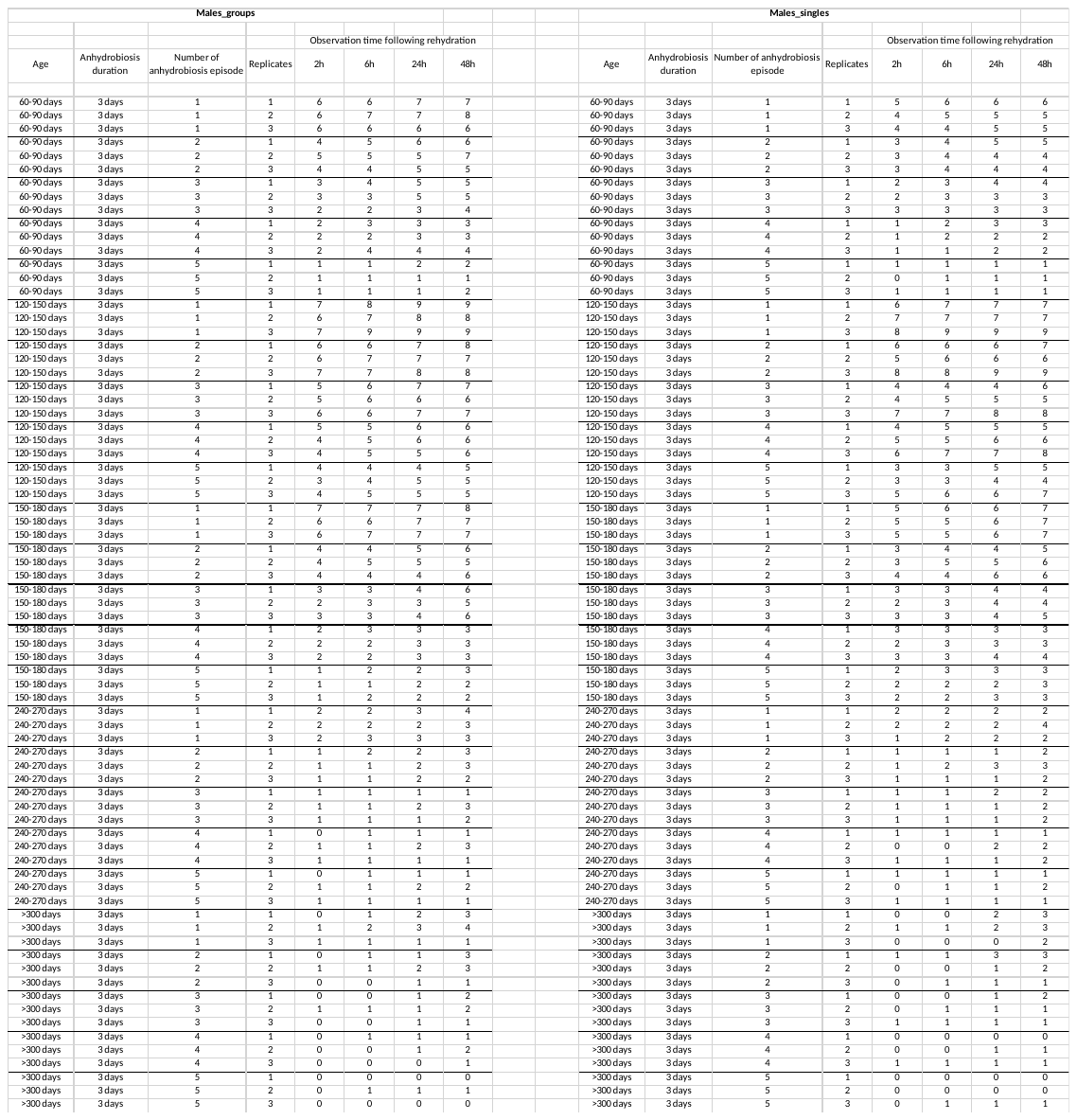

Females: two episodes of long anhydrobiosis

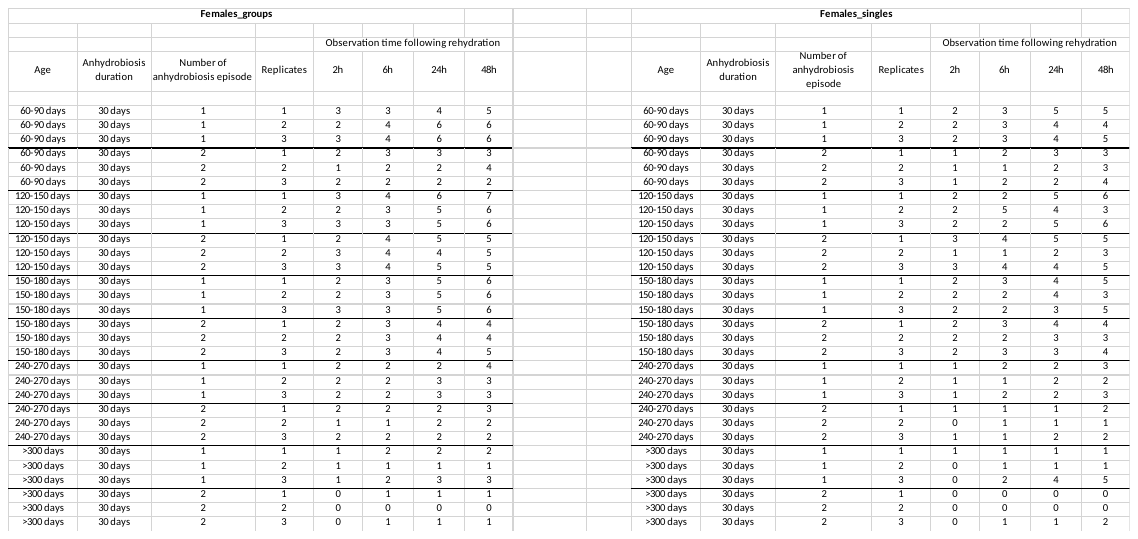

Males: two episodes of long anhydrobiosis

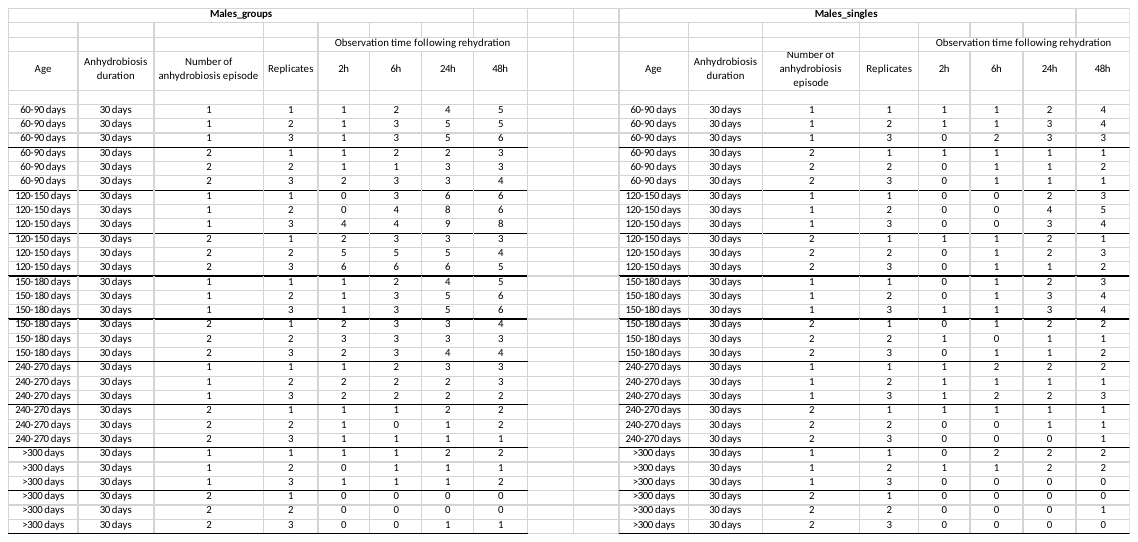

*Statistical analysis – assumptions for ANOVA*

Data were tested for normal distribution using Kolmogorov-Smirnov (KS) tests [5] and Levene's tests for homogeneity of group variances [6]. Left skewed distributions and non-homogeneity of variances were observed for some variables. Although, in general, the Type I error and the power of the F-statistic are not altered by violations of normality, as predicted by the central limit theorem, especially for large sample sizes [e.g., 7], ANOVAs were performed using the square root transformed reflected variable, x(transf.) = sqrt{1+max(x)-x}. This data transformation gave the best results in the context of the tests. One variable, the number of active individuals 2 h after rehydration, was not normally distributed, d = 0.134, p<0.05).

5. Lilliefors, H. W. On the Kolmogorov-Smirnov Test for normality with mean and variance unknown. *J. Am. Stat. Assoc.* **62,** 399–402. <https://doi.org/10.1080/01621459.1967.10482916> (1967).
6. Levene, H. Robust tests for equality of variances in *Contributions to probability and statistics* (eds. Olkin, I., Ghurye, S. G., Hoeffding, W., Madow, W. G. & Mann, H. B.) 278–292 (Stanford University Press: Stanford) (1960).
7. Keselman, J. C., Lix, L. M., & Keselman, H. J. The analysis of repeated measurements: A quantitative research synthesis. *Br. J. Math. Stat. Psychol.* **49,** 275–298. <https://doi.org/10.1111/j.2044-8317.1996.tb01089> (1996).

**Table S3.** Multivariate repeated measures ANOVA factor analysis for combination of the number and duration of anhydrobiosis episodes (anhydrobiosis duration), the presence of other individuals (i.e., group influence), age and sex of individuals (as independent variables) on the number of active individuals of *P. experimentalis* observed at 2h-48h after rehydration. ANOVA was performed using the square root transformed reflected variable, x(transf.) = sqrt{1+max(x)-x}; the test unit was 30 individuals (3 replicates x 10 individuals). in red - statistically significant results. SS - Sum of Squares, MS - Mean Squares, F - F ratio and P - *p* values are given.

| **Effect** | **SS** | **Degree of freedom** | **MS** | **F** | **P** |
| --- | --- | --- | --- | --- | --- |
| Intercept | 3750,842 | 1 | 3750,842 | 2361,121 | 0,0000000 |
| {1} anhydrobiosis duration | 24,362 | 1 | 24,362 | 15,336 | 0,0001646 |
| {2} group influence | 18,285 | 1 | 18,285 | 11,511 | 0,0009928 |
| {3} sex | 7,159 | 1 | 7,159 | 4,507 | 0,0362302 |
| {4} age | 196,546 | 4 | 49,136 | 30,931 | 0,0000000 |
| anhydrobiosis duration*group influence | 4,488 | 1 | 4,488 | 2,825 | 0,0959270 |
| anhydrobiosis duration*sex | 2,631 | 1 | 2,631 | 1,656 | 0,2010482 |
| group influence*sex | 2,736 | 1 | 2,736 | 1,722 | 0,1924293 |
| anhydrobiosis duration*age | 10,134 | 4 | 2,533 | 1,595 | 0,1815664 |
| group influence*age | 0,791 | 4 | 0,198 | 0,124 | 0,9733427 |
| sex*age | 0,920 | 4 | 0,230 | 0,145 | 0,9648783 |
| anhydrobiosis duration*group influence*sex | 2,514 | 1 | 2,514 | 1,582 | 0,2113643 |
| anhydrobiosis duration*group influence*age | 4,241 | 4 | 1,060 | 0,667 | 0,6161241 |
| anhydrobiosis duration*sex*age | 3,140 | 4 | 0,785 | 0,494 | 0,7400715 |
| group influence*sex*age | 2,060 | 4 | 0,515 | 0,324 | 0,8611674 |
| anhydrobiosis duration*group influence*sex*age | 1,080 | 4 | 0,270 | 0,170 | 0,9532145 |
| Error | 158,859 | 100 | 1,589 |  |  |
| {5}OBS.TIME | 28,290 | 3 | 9,430 | 172,376 | 0,0000000 |
| OBS.TIME*anhydrobiosis duration | 2,182 | 3 | 0,727 | 13,298 | 0,0000000 |
| OBS.TIME*group influence | 0,473 | 3 | 0,158 | 2,883 | 0,0360823 |
| OBS.TIME*sex | 0,209 | 3 | 0,070 | 1,271 | 0,2844629 |
| OBS.TIME*age | 2,305 | 12 | 0,192 | 3,511 | 0,0000698 |
| OBS.TIME*anhydrobiosis duration*group influence | 0,441 | 3 | 0,147 | 2,687 | 0,0466582 |
| OBS.TIME*anhydrobiosis duration*sex | 0,329 | 3 | 0,110 | 2,004 | 0,1134477 |
| OBS.TIME*group influence*sex | 0,107 | 3 | 0,036 | 0,650 | 0,5833932 |
| OBS.TIME*anhydrobiosis duration*age | 5,761 | 12 | 0,480 | 8,776 | 0,0000000 |
| OBS.TIME*group influence*age | 1,067 | 12 | 0,089 | 1,625 | 0,0837031 |
| OBS.TIME*sex*age | 1,665 | 12 | 0,139 | 2,537 | 0,0033537 |
| 5*1*2*3 | 0,466 | 3 | 0,155 | 2,837 | 0,0383341 |
| 5*1*2*4 | 0,765 | 12 | 0,064 | 1,166 | 0,3072984 |
| 5*1*3*4 | 1,712 | 12 | 0,143 | 2,608 | 0,0025540 |
| 5*2*3*4 | 0,682 | 12 | 0,057 | 1,039 | 0,4131341 |
| 5*1*2*3*4 | 0,964 | 12 | 0,080 | 1,468 | 0,1351782 |
| Error | 16,412 | 300 | 0,055 |  |  |

**Table S4.** Number of active individuals of *P. experimentalis* observed at 2h-48h after rehydration, with an emphasis on their age and duration of anhydrobiosis ((long anhydrobiosis - two episodes lasting 30 days each; short anhydrobiosis – five episodes lasting 3 days each). Based on the square root transformed reflected variable, x(transf.) = sqrt{1+max(x)-x}. Mean, SE - Standard Error, and 95% confidence interval for the population mean are given.

| **Subclass** | **anhydrobiosis** | **age class** | **OBS.TIME** | **Mean** | **SE** | **-95%** | **95%** | **N** |
| --- | --- | --- | --- | --- | --- | --- | --- | --- |
| 1 | long | 1 | 2h | 2,385 | 0,232 | 1,925 | 2,846 | 8 |
| 2 | long | 1 | 6h | 2,723 | 0,224 | 2,278 | 3,169 | 8 |
| 3 | long | 1 | 24h | 3,174 | 0,229 | 2,719 | 3,629 | 8 |
| 4 | long | 1 | 48h | 3,313 | 0,250 | 2,818 | 3,808 | 8 |
| 5 | long | 2 | 2h | 2,483 | 0,232 | 2,023 | 2,944 | 8 |
| 6 | long | 2 | 6h | 2,948 | 0,224 | 2,503 | 3,393 | 8 |
| 7 | long | 2 | 24h | 3,715 | 0,229 | 3,260 | 4,170 | 8 |
| 8 | long | 2 | 48h | 3,695 | 0,250 | 3,200 | 4,190 | 8 |
| 9 | long | 3 | 2h | 2,303 | 0,232 | 1,842 | 2,763 | 8 |
| 10 | long | 3 | 6h | 2,756 | 0,224 | 2,311 | 3,201 | 8 |
| 11 | long | 3 | 24h | 3,351 | 0,229 | 2,896 | 3,807 | 8 |
| 12 | long | 3 | 48h | 3,470 | 0,250 | 2,975 | 3,965 | 8 |
| 13 | long | 4 | 2h | 2,086 | 0,232 | 1,626 | 2,547 | 8 |
| 14 | long | 4 | 6h | 2,223 | 0,224 | 1,778 | 2,669 | 8 |
| 15 | long | 4 | 24h | 2,472 | 0,229 | 2,017 | 2,927 | 8 |
| 16 | long | 4 | 48h | 2,515 | 0,250 | 2,020 | 3,010 | 8 |
| 17 | long | 5 | 2h | 1,695 | 0,232 | 1,235 | 2,155 | 8 |
| 18 | long | 5 | 6h | 1,729 | 0,224 | 1,284 | 2,174 | 8 |
| 19 | long | 5 | 24h | 1,864 | 0,229 | 1,409 | 2,319 | 8 |
| 20 | long | 5 | 48h | 1,770 | 0,250 | 1,275 | 2,265 | 8 |
| 21 | short | 1 | 2h | 3,093 | 0,147 | 2,802 | 3,384 | 20 |
| 22 | short | 1 | 6h | 3,336 | 0,142 | 3,054 | 3,618 | 20 |
| 23 | short | 1 | 24h | 3,542 | 0,145 | 3,255 | 3,830 | 20 |
| 24 | short | 1 | 48h | 3,440 | 0,158 | 3,127 | 3,754 | 20 |
| 25 | short | 2 | 2h | 3,978 | 0,147 | 3,686 | 4,269 | 20 |
| 26 | short | 2 | 6h | 4,151 | 0,142 | 3,870 | 4,433 | 20 |
| 27 | short | 2 | 24h | 4,331 | 0,145 | 4,043 | 4,619 | 20 |
| 28 | short | 2 | 48h | 4,252 | 0,158 | 3,939 | 4,566 | 20 |
| 29 | short | 3 | 2h | 3,285 | 0,147 | 2,994 | 3,577 | 20 |
| 30 | short | 3 | 6h | 3,471 | 0,142 | 3,189 | 3,752 | 20 |
| 31 | short | 3 | 24h | 3,658 | 0,145 | 3,370 | 3,946 | 20 |
| 32 | short | 3 | 48h | 3,704 | 0,158 | 3,391 | 4,017 | 20 |
| 33 | short | 4 | 2h | 2,215 | 0,147 | 1,924 | 2,506 | 20 |
| 34 | short | 4 | 6h | 2,389 | 0,142 | 2,107 | 2,671 | 20 |
| 35 | short | 4 | 24h | 2,608 | 0,145 | 2,320 | 2,896 | 20 |
| 36 | short | 4 | 48h | 2,721 | 0,158 | 2,408 | 3,034 | 20 |
| 37 | short | 5 | 2h | 1,556 | 0,147 | 1,265 | 1,848 | 20 |
| 38 | short | 5 | 6h | 1,827 | 0,142 | 1,546 | 2,109 | 20 |
| 39 | short | 5 | 24h | 2,134 | 0,145 | 1,846 | 2,422 | 20 |
| 40 | short | 5 | 48h | 2,212 | 0,158 | 1,899 | 2,525 | 20 |
| class |  |  |  |  |  |  |  |  |
| 1 | long | all | 2h-48h | 2,634 | 0,100 | 2,436 | 2,831 | 40 |
| 2 | short | all | 2h-48h | 3,095 | 0,063 | 2,970 | 3,220 | 100 |

**Table S5.** Correlations between observation time after rehydration and duration of anhydrobiosis (long vs. short). Anhydrobiosis duration denotes combination of the number and duration of anhydrobiosis episodes. Correlation coefficients in red are significant at *p*< 0.05.

| **Obs. time after rehydration** | **2h** | **6h** | **24h** | **48h** | **anhydrobiosis duration** |
| --- | --- | --- | --- | --- | --- |
| 2h | 1,000 | 0,939 | 0,887 | 0,857 | 0,280 |
| 6h | 0,939 | 1,000 | 0,960 | 0,926 | 0,254 |
| 24h | 0,887 | 0,960 | 1,000 | 0,979 | 0,157 |
| 48h | 0,857 | 0,926 | 0,979 | 1,000 | 0,145 |
| anhydrobiosis duration | 0,280 | 0,254 | 0,157 | 0,145 | 1,000 |

**Table S6.** Number of active individuals after rehydration (2h - 48h) considering the first two episodes of short (SA) and long anhydrobiosis (LA); Mean, SE - standard error, and 95% confidence interval are given.

|  | **episodes** | **Mean_2h** | **SE_2h** | **-95%_6h** | **95%_2h** | **Mean_6h** | **SE_6h** | **-95%_6h** | **95%_6h** | **Mean_24h** | **SE_24h** | **-95%_24h** | **95%_24h** | **Mean_48h** | **SE_48h** | **-95%_48h** | **95%_48h** | **N** |
| --- | --- | --- | --- | --- | --- | --- | --- | --- | --- | --- | --- | --- | --- | --- | --- | --- | --- | --- |
| 1 | LA_E1 | 2,220 | 0,213 | 1,795 | 2,645 | 2,634 | 0,201 | 2,232 | 3,035 | 3,291 | 0,197 | 2,898 | 3,684 | 3,362 | 0,183 | 2,997 | 3,727 | 20 |
| 2 | LA_E2 | 2,161 | 0,213 | 1,736 | 2,586 | 2,318 | 0,201 | 1,917 | 2,719 | 2,539 | 0,197 | 2,146 | 2,932 | 2,543 | 0,183 | 2,179 | 2,908 | 20 |
| 3 | SA_E1 | 3,586 | 0,213 | 3,161 | 4,011 | 3,803 | 0,201 | 3,402 | 4,204 | 4,023 | 0,197 | 3,630 | 4,417 | 4,073 | 0,183 | 3,709 | 4,438 | 20 |
| 4 | SA_E2 | 3,215 | 0,213 | 2,790 | 3,640 | 3,456 | 0,201 | 3,054 | 3,857 | 3,657 | 0,197 | 3,264 | 4,050 | 3,767 | 0,183 | 3,402 | 4,131 | 20 |

**Table S7.** Number of active individuals for groups and single animals by the duration of anhydrobiosis (combination of the number and duration of anhydrobiosis episodes) and observation time (Obs. time) after rehydration (2h-48h); Mean, SE - Standard Error, and 95% confidence interval for the population mean are given; LA - long anhydrobiosis; SA - short anhydrobiosis.

| **#** | **anhydrobiosis** | **in groups or single animals** | **Obs. time** | **Mean** | **SE** | **-95%** | **95%** | **N** |
| --- | --- | --- | --- | --- | --- | --- | --- | --- |
| 1 | LA | group | 2h | 2,592 | 0,147 | 2,301 | 2,884 | 20 |
| 2 | LA | group | 6h | 2,767 | 0,142 | 2,485 | 3,049 | 20 |
| 3 | LA | group | 24h | 3,186 | 0,145 | 2,898 | 3,473 | 20 |
| 4 | LA | group | 48h | 3,186 | 0,158 | 2,873 | 3,499 | 20 |
| 5 | LA | single | 2h | 1,789 | 0,147 | 1,498 | 2,080 | 20 |
| 6 | LA | single | 6h | 2,185 | 0,142 | 1,903 | 2,466 | 20 |
| 7 | LA | single | 24h | 2,645 | 0,145 | 2,357 | 2,933 | 20 |
| 8 | LA | single | 48h | 2,720 | 0,158 | 2,407 | 3,033 | 20 |
| 9 | SA | group | 2h | 2,932 | 0,093 | 2,748 | 3,116 | 50 |
| 10 | SA | group | 6h | 3,129 | 0,090 | 2,951 | 3,307 | 50 |
| 11 | SA | group | 24h | 3,349 | 0,092 | 3,167 | 3,531 | 50 |
| 12 | SA | group | 48h | 3,375 | 0,100 | 3,177 | 3,573 | 50 |
| 13 | SA | single | 2h | 2,719 | 0,093 | 2,535 | 2,903 | 50 |
| 14 | SA | single | 6h | 2,941 | 0,090 | 2,763 | 3,119 | 50 |
| 15 | SA | single | 24h | 3,160 | 0,092 | 2,978 | 3,342 | 50 |
| 16 | SA | single | 48h | 3,157 | 0,100 | 2,959 | 3,355 | 50 |

**Table S8.** Number of active females (F) and males (M) by the duration of anhydrobiosis (combination of the number and duration of anhydrobiosis episodes) and observation time (Obs. Time) after rehydration (2h-48h). Mean, SE - Standard Error, and 95% for the population mean are given .

| **#** | **age** | **anhydrobiosis duration** | **sex** | **Obs. Time** | **Mean** | **SE** | **-95%** | **95%** | **N** |
| --- | --- | --- | --- | --- | --- | --- | --- | --- | --- |
| 1 | 1 | LA | M | 2h | 2,247 | 0,328 | 1,596 | 2,898 | 4 |
| 2 | 1 | LA | M | 6h | 2,470 | 0,317 | 1,841 | 3,100 | 4 |
| 3 | 1 | LA | M | 24h | 2,968 | 0,324 | 2,325 | 3,612 | 4 |
| 4 | 1 | LA | M | 48h | 3,120 | 0,353 | 2,420 | 3,820 | 4 |
| 5 | 1 | LA | F | 2h | 2,524 | 0,328 | 1,873 | 3,175 | 4 |
| 6 | 1 | LA | F | 6h | 2,976 | 0,317 | 2,346 | 3,606 | 4 |
| 7 | 1 | LA | F | 24h | 3,380 | 0,324 | 2,737 | 4,024 | 4 |
| 8 | 1 | LA | F | 48h | 3,507 | 0,353 | 2,807 | 4,207 | 4 |
| 9 | 1 | SA | M | 2h | 2,921 | 0,208 | 2,509 | 3,333 | 10 |
| 10 | 1 | SA | M | 6h | 3,189 | 0,201 | 2,790 | 3,587 | 10 |
| 11 | 1 | SA | M | 24h | 3,401 | 0,205 | 2,994 | 3,808 | 10 |
| 12 | 1 | SA | M | 48h | 3,311 | 0,223 | 2,868 | 3,754 | 10 |
| 13 | 1 | SA | F | 2h | 3,265 | 0,208 | 2,853 | 3,676 | 10 |
| 14 | 1 | SA | F | 6h | 3,483 | 0,201 | 3,085 | 3,882 | 10 |
| 15 | 1 | SA | F | 24h | 3,684 | 0,205 | 3,277 | 4,091 | 10 |
| 16 | 1 | SA | F | 48h | 3,570 | 0,223 | 3,127 | 4,013 | 10 |
| 17 | 2 | LA | M | 2h | 2,098 | 0,328 | 1,447 | 2,749 | 4 |
| 18 | 2 | LA | M | 6h | 2,584 | 0,317 | 1,954 | 3,214 | 4 |
| 19 | 2 | LA | M | 24h | 3,596 | 0,324 | 2,952 | 4,239 | 4 |
| 20 | 2 | LA | M | 48h | 3,462 | 0,353 | 2,762 | 4,163 | 4 |
| 21 | 2 | LA | F | 2h | 2,869 | 0,328 | 2,218 | 3,520 | 4 |
| 22 | 2 | LA | F | 6h | 3,312 | 0,317 | 2,682 | 3,942 | 4 |
| 23 | 2 | LA | F | 24h | 3,833 | 0,324 | 3,190 | 4,477 | 4 |
| 24 | 2 | LA | F | 48h | 3,928 | 0,353 | 3,228 | 4,628 | 4 |
| 25 | 2 | SA | M | 2h | 4,101 | 0,208 | 3,689 | 4,513 | 10 |
| 26 | 2 | SA | M | 6h | 4,314 | 0,201 | 3,915 | 4,712 | 10 |
| 27 | 2 | SA | M | 24h | 4,488 | 0,205 | 4,081 | 4,895 | 10 |
| 28 | 2 | SA | M | 48h | 4,469 | 0,223 | 4,026 | 4,911 | 10 |
| 29 | 2 | SA | F | 2h | 3,854 | 0,208 | 3,443 | 4,266 | 10 |
| 30 | 2 | SA | F | 6h | 3,989 | 0,201 | 3,590 | 4,387 | 10 |
| 31 | 2 | SA | F | 24h | 4,174 | 0,205 | 3,767 | 4,581 | 10 |
| 32 | 2 | SA | F | 48h | 4,036 | 0,223 | 3,593 | 4,479 | 10 |
| 33 | 3 | LA | M | 2h | 1,914 | 0,328 | 1,263 | 2,565 | 4 |
| 34 | 3 | LA | M | 6h | 2,474 | 0,317 | 1,844 | 3,103 | 4 |
| 35 | 3 | LA | M | 24h | 3,106 | 0,324 | 2,463 | 3,750 | 4 |
| 36 | 3 | LA | M | 48h | 3,248 | 0,353 | 2,548 | 3,948 | 4 |
| 37 | 3 | LA | F | 2h | 2,691 | 0,328 | 2,040 | 3,342 | 4 |
| 38 | 3 | LA | F | 6h | 3,038 | 0,317 | 2,408 | 3,668 | 4 |
| 39 | 3 | LA | F | 24h | 3,597 | 0,324 | 2,953 | 4,240 | 4 |
| 40 | 3 | LA | F | 48h | 3,693 | 0,353 | 2,993 | 4,393 | 4 |
| 41 | 3 | SA | M | 2h | 3,169 | 0,208 | 2,757 | 3,580 | 10 |
| 42 | 3 | SA | M | 6h | 3,378 | 0,201 | 2,980 | 3,777 | 10 |
| 43 | 3 | SA | M | 24h | 3,612 | 0,205 | 3,205 | 4,019 | 10 |
| 44 | 3 | SA | M | 48h | 3,706 | 0,223 | 3,263 | 4,148 | 10 |
| 45 | 3 | SA | F | 2h | 3,402 | 0,208 | 2,990 | 3,814 | 10 |
| 46 | 3 | SA | F | 6h | 3,563 | 0,201 | 3,165 | 3,962 | 10 |
| 47 | 3 | SA | F | 24h | 3,704 | 0,205 | 3,297 | 4,111 | 10 |
| 48 | 3 | SA | F | 48h | 3,702 | 0,223 | 3,259 | 4,145 | 10 |
| 49 | 4 | LA | M | 2h | 1,966 | 0,328 | 1,315 | 2,617 | 4 |
| 50 | 4 | LA | M | 6h | 2,060 | 0,317 | 1,431 | 2,690 | 4 |
| 51 | 4 | LA | M | 24h | 2,312 | 0,324 | 1,668 | 2,955 | 4 |
| 52 | 4 | LA | M | 48h | 2,312 | 0,353 | 1,611 | 3,012 | 4 |
| 53 | 4 | LA | F | 2h | 2,207 | 0,328 | 1,556 | 2,858 | 4 |
| 54 | 4 | LA | F | 6h | 2,386 | 0,317 | 1,756 | 3,016 | 4 |
| 55 | 4 | LA | F | 24h | 2,632 | 0,324 | 1,988 | 3,275 | 4 |
| 56 | 4 | LA | F | 48h | 2,718 | 0,353 | 2,018 | 3,418 | 4 |
| 57 | 4 | SA | M | 2h | 2,163 | 0,208 | 1,751 | 2,575 | 10 |
| 58 | 4 | SA | M | 6h | 2,318 | 0,201 | 1,920 | 2,717 | 10 |
| 59 | 4 | SA | M | 24h | 2,515 | 0,205 | 2,108 | 2,922 | 10 |
| 60 | 4 | SA | M | 48h | 2,613 | 0,223 | 2,170 | 3,056 | 10 |
| 61 | 4 | SA | F | 2h | 2,267 | 0,208 | 1,856 | 2,679 | 10 |
| 62 | 4 | SA | F | 6h | 2,460 | 0,201 | 2,061 | 2,858 | 10 |
| 63 | 4 | SA | F | 24h | 2,701 | 0,205 | 2,294 | 3,108 | 10 |
| 64 | 4 | SA | F | 48h | 2,829 | 0,223 | 2,387 | 3,272 | 10 |
| 65 | 5 | LA | M | 2h | 2,037 | 0,328 | 1,386 | 2,688 | 4 |
| 66 | 5 | LA | M | 6h | 1,500 | 0,317 | 0,870 | 2,130 | 4 |
| 67 | 5 | LA | M | 24h | 1,618 | 0,324 | 0,975 | 2,262 | 4 |
| 68 | 5 | LA | M | 48h | 1,559 | 0,353 | 0,859 | 2,259 | 4 |
| 69 | 5 | LA | F | 2h | 1,354 | 0,328 | 0,703 | 2,005 | 4 |
| 70 | 5 | LA | F | 6h | 1,958 | 0,317 | 1,328 | 2,588 | 4 |
| 71 | 5 | LA | F | 24h | 2,109 | 0,324 | 1,466 | 2,753 | 4 |
| 72 | 5 | LA | F | 48h | 1,981 | 0,353 | 1,281 | 2,681 | 4 |
| 73 | 5 | SA | M | 2h | 1,395 | 0,208 | 0,983 | 1,807 | 10 |
| 74 | 5 | SA | M | 6h | 1,650 | 0,201 | 1,252 | 2,049 | 10 |
| 75 | 5 | SA | M | 24h | 2,068 | 0,205 | 1,661 | 2,475 | 10 |
| 76 | 5 | SA | M | 48h | 2,140 | 0,223 | 1,697 | 2,583 | 10 |
| 77 | 5 | SA | F | 2h | 1,718 | 0,208 | 1,306 | 2,130 | 10 |
| 78 | 5 | SA | F | 6h | 2,004 | 0,201 | 1,606 | 2,403 | 10 |
| 79 | 5 | SA | F | 24h | 2,200 | 0,205 | 1,793 | 2,607 | 10 |
| 80 | 5 | SA | F | 48h | 2,284 | 0,223 | 1,842 | 2,727 | 10 |

**Figure S1**. *Paramacrobiotus experimentalis* individuals in a group during the tun state. The young females (age group 2) are shown during the first short anhydrobiosis episode; black arrows indicate the tuns.

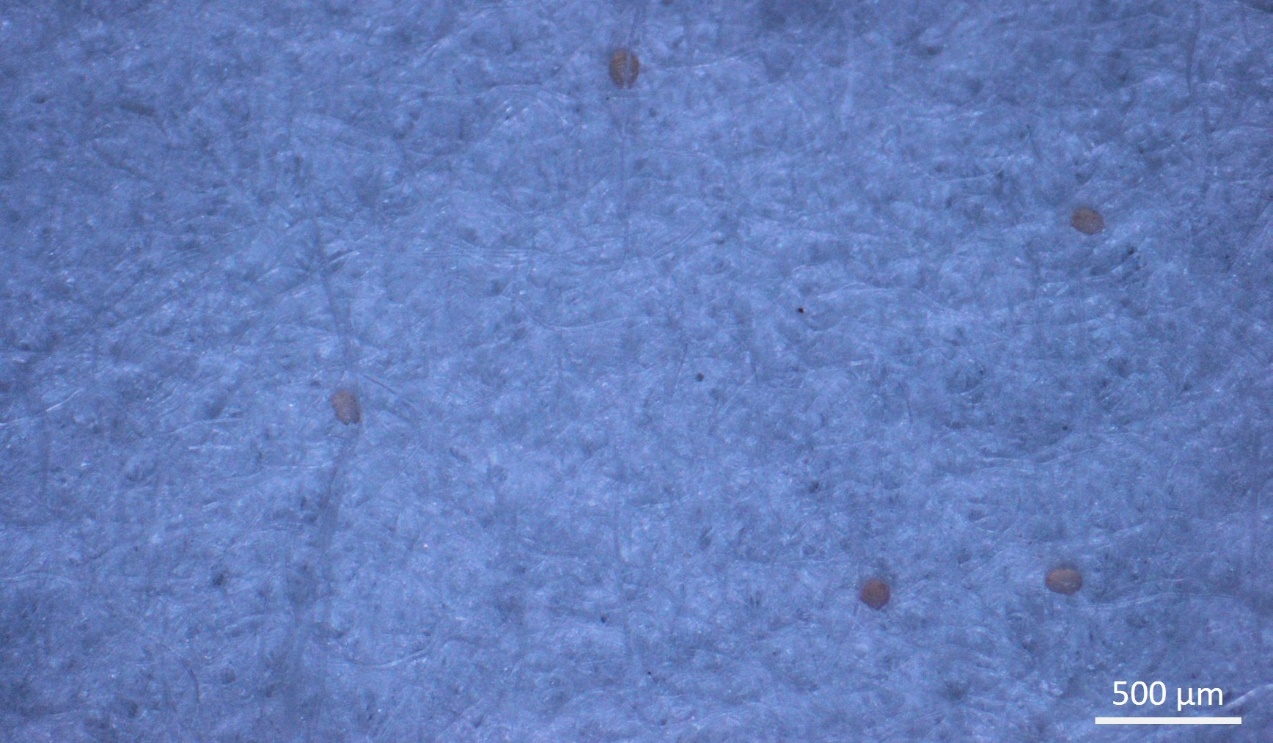
